# Loss of Paneth cells dysregulates gut ILC subsets and enhances weight gain response to high fat diet in a mouse model

**DOI:** 10.1101/2024.03.29.587349

**Authors:** Marisa R. Joldrichsen, Eunsoo Kim, Haley E. Steiner, Yea Ji Jeong, Christopher Premanandan, Willa Hsueh, Ouliana Ziouzenkova, Estelle Cormet-Boyaka, Prosper N. Boyaka

**Affiliations:** Department of Veterinary Biosciences, The Ohio State University, Columbus, OH; Department of Internal Medicine, The Ohio State University, Columbus, OH; Department of Human Sciences, The Ohio State University, Columbus, OH; College of Pharmacy, Pusan National University, Busan, Republic of Korea

**Keywords:** Paneth cells, Innate lymphoid cells, obesity, intestine

## Abstract

Obesity has been associated with dysbiosis, but innate mechanisms linking intestinal epithelial cell subsets and obesity remain poorly understood. Using mice lacking Paneth cells (Sox9^ΔIEC^ mice), small intestinal epithelial cells specialized in the production of antimicrobial products and cytokines, we show that dysbiosis alone does not induce obesity or metabolic disorders. Loss of Paneth cells reduced ILC3 and increased ILC2 numbers in the intestinal lamina propria. High-fat diet (HFD) induced higher weight gain and more severe metabolic disorders in Sox9^ΔIEC^ mice. Further, HFD enhances the number of ILC1 in the intestinal lamina propria of Sox9^ΔIEC^ mice and increases intestinal permeability and the accumulation of immune cells (inflammatory macrophages and T cells, and B cells) in abdominal fat tissues of obese Sox9^ΔIEC^. Transplantation of fecal materials from Sox9^ΔIEC^ mice in germ-free mice before HFD further confirmed the regulatory role of Paneth cells for gut ILC subsets and the development of obesity.

## Introduction

Obesity is a growing epidemic across the world and in the United States alone it is currently estimated that at least 1/3 of United States adults or roughly 109.8 million people are considered obese [1–5]. The rise in obesity has been linked to lifestyle changes, including the consumption of processed food, increased calories intake, and a decrease in physical activity [6–8]. With the incidence of obesity only expected to increase in the coming years, it has become crucial to understand the cause and identify potential treatments to combat this growing epidemic [6–8]. Alteration of gut microbial community (dysbiosis) have been shown to be a driver of obesity [9]. The small and large intestine of the gastrointestinal system are major connection point between the outside world (foodstuffs, nutrients, microbes, etc.), and the cells of human body [10]. Thus, intestinal epithelial cells play an integral role in the absorption of nutrients and regulation of interactions between the microbiome and host immune cells [11]. While a balanced microbiome is important for gastrointestinal (GI) health, and dysbiosis alters several biological processes throughout the host, the role of individual epithelial cell subsets in the development of obesity is poorly understood.

Intestinal epithelial cells are the first line of host cells in contact with commensal microbes, and pathogens that enter the body through the GI system [10,11]. Goblet cells produce mucus, which limits access of most microbes to the epithelial layer and regulates entry of luminal content via the goblet cell associated passages [12]. Enterocytes represent the major subset of intestinal epithelial cells and as such are the major sensors of microbes in the GI tract. They are also regulator of gut immune homeostasis, and initiation/stimulation of inflammatory responses and adaptive immunity [13]. Commensal microbe sensing and activation of antimicrobial responses can lead to dysbiosis [14–16] and a variety of immunological and metabolic disease [17,18], including obesity [19]. Paneth cells are found in the crypts of the small intestine. It has been shown that Paneth cells directly sense gut commensals and maintain homeostasis at the intestinal host-microbial interface through production of anti-microbials and cytokines [20,21]. Thus, Paneth cells produce a-defensins, lysozyme, and phospholipase A_2_, and contribute to immune homeostasis through production of cytokines (TNF-α and IL-17A) [22–25]. As a main source of anti-microbial products in the gut, Paneth cells are believed to play a protective role in the development of metabolic disorders and obesity [20,26–28]. It was previously reported that unfolded protein responses limit secretion of HD5 and lysozyme by Paneth cells of severely obese individuals [29]. Furthermore, western diet was shown to promote Paneth cell dysfunction through mechanisms dependent on farnesoid X receptor (FXR) and type I interferon signaling [30]. However, it remains unclear how Paneth cells dysfunction regulate innate responses which control the development of metabolic disorders and weight gain [31].

Using mice which lack Paneth cells [32–34], we examined the role of Paneth cells on homeostatic regulation of mucosal immune cells of the GI tract and after exposure to a western or high fat diet (HFD). Here we report that lack of Paneth cells switches the profile of ILCs in the intestinal lamina propria from a mostly ILC3 to an ILC2 environment. We also report that without Paneth cells, HFD more effectively increases weight gain and metabolic disorders as indicated by increased glucose intolerance. The fecal material of mice lacking Paneth cells could transfer the hyper response to HFD phenotype to recipient germ-free mice indicating that that Paneth cells play role in the protection again metabolic disorders and obesity through regulation of ILC subsets in gut tissues.

## Materials and Methods

### Mice

C57BL/6 purchased from The Jackson Laboratory (Bay Harbor, ME, and Columbus, OH) and Sox9^ΔIEC^ mice obtained from Dr. Mori-Akiyama (The University of Texas, M.D. Anderson Cancer Center). Mice were co-housed with in-house in-bred wild-type C57BL/6 and Cre-negative controls, maintained under a specific pathogen-free environment, fed the same standard mouse chow, and provided 12:12-hour light and dark cycle in our animal facility at The Ohio State University. Germ-Free mice were obtained from Charles River Laboratories (Wilmington, MA) and maintained in germ-free isolators in our germ-free facility. Food and all supplies for germ-free mice were autoclaved and sterilized before entry into the isolators, and isolators were tested monthly for bacterial contamination. For use in experiments, germ-free mice were transferred into sterile Allentown cages, using the Allentown rack and hood system (Allentown, NJ). All mice were given food and water *ad libitum* unless short-term fasting (4-8 hours) was needed for an experiment. Mice were anesthetized by injection of Ketamine and euthanized exposure to CO_2_ followed by cervical dissociation. All protocols were reviewed and approved by the institutional Animal Care and Use Committee (IACUC) of the Ohio State University and followed the federal and ARRIVE guidelines to avoid unnecessary pain and distress and to minimize animal suffering.

### Diet-induced obesity

Male mice aged 8-12 weeks were given a high fat diet (HFD, 60% kcal fat, Envigo Teklad Diets, Madison, WI) *ad libitum* over thirteen weeks. Body weight was recorded weekly for 13 weeks. Control animals were fed ad libitum a standard mouse chow which contained the same basic nutrients with the exception that it only provides 10% of total calories from fat.

### Intraperitoneal glucose tolerance test (ipGTT)

To examine the ability of mice under HDF to regulate serum glucose levels, ipGTT were performed on weeks 12 and/or 13 of the treatment. Thus, mice were starved for 8 hours, and basal blood glucose levels were measured using the Contour Next EZ glucose meter and strips (Bayer, Germany). Mice were then intraperitoneally injected a sugar solution (33% dextrose administered at 1g/kg of body weight), and their blood glucose levels were measured every 15 minutes for 120 min.

### Evaluation of intestinal permeability

For evaluation of intestinal permeability, mice were deprived of water for 4hrs and orally administered 200 ml of PBS containing 8 mg of FITC-dextran (4000 Da; FD4, Sigma Aldrich, Saint Louis, MO) per 20 g of body weight. After an additional 2 hours blood was collected and serum FITC-dextran levels were measured with a Victor V multiparameter plate reader (Perkin Elmer, Waltham, MA). Quantification of serum FITC-dextran levels was performed with the WorkOut 2.5 software by extrapolation against know concentrations of FITC-dextran standard.

### 16s RNA analysis of gut microbial community

Freshly emitted fecal pellets were collected, immediately placed on dry ice, and stored at −80°C. Bacterial DNA was extracted by conventional methods (Qiagen, Valencia, CA, USA) and bacterial communities were analyzed by bacterial tag encoded FLX amplicon pyrosequencing (Roche, Branford, CT, USA) as previously described [35,36]. Linear discriminant analysis (LDA) was performed using the Galaxy web application [Huttenhower lab, Harvard university (https://huttenhower.sph.harvard.edu/galaxy/)] and the threshold was set at 2.0 [36,37]. Principal component analysis was done at the phylum level using Prism, with the eigen values determining the scale for the graph. Eigen values were selected according to the Kaiser rule with values being over 1.0. The data were visualized as score plots with each group being a different color.

### Fecal material transplants

Freshly emitted fecal pellets were collected and immediately placed on ice. Fecal material content was then normalized by dilution in sterile PBS (1mg per 0.1g of fecal material) and vortexed thoroughly. Particles were removed by filtration through 70 μm strainers and mice were given 300ml of solution by gavage.

### Isolation of abdominal fat cells

Abdominal tissues were collected at euthanasia and immediately placed in cold sterile PBS on ice. Adipose tissue cells were then isolated by a modification of a previously reported protocol by others [38]. Briefly, the tissues were placed in a collagenase digestion solution (PBS, with 0.5% BSA, 10 mM CaCl_2_, and 4 mg/ml collagenase) and incubated in a rotational shaker for 20 minutes at 37°C. The homogenates were then filtered through 70 mm cell strainers and spun down at 500 x g for 10 minutes at 4°C. Erythrocytes were lysed by incubation in ACK buffer followed by washes in complete RPMI, and the pellets resuspended in FACS buffer for cell counting and flow cytometry analysis.

### Isolation of lamina propria cells

Small intestinal tissues were collected at euthanasia and immediately flushed with cold PBS to remove fecal materials. After removal of fat and visible Peyer’s Patches, the small intestine tissues were opened longitudinally, and cut into 1-2 cm pieces which were placed in a pre-digestion buffer (1x PBS,10mM Hepes, 5mM EDTA, and 5% FCS). The tissues were incubated in a rotational shaker for 30 minutes at 37°C, supernatants were removed for collection of intraepithelial lymphoid cells (IEL), and the procedure was repeated 3 times. The remaining tissues were then placed in a digestion buffer (PBS with 10mM Hepes, 0.5mg/ml collagenase, and 5% FCS) and incubated in a rotational shaker for 30 minutes at 37°C for isolation of cells from the lamina propria. Cells isolated from the small intestinal intraepithelial compartment and lamina propria were further purified using a 4-layers Percoll gradient (75%, 60%, 40%, and 20%). The cells at the interface of the 40% and 60% Percoll layers were collected and washed with complete RPMI before counting and flow cytometry analysis.

### Isolation of cells from mesenteric lymph nodes

Mesenteric lymph nodes were collected and immediately placed in RPMI on ice. The tissues were mechanically dissociated by passage through 70 mm cell strainers and cells washed 3 times with complete RPMI before counting and flow cytometry analysis.

### Flow cytometry analysis of immune cells

Immune cell subsets in fat tissues, intestinal tissues (IEL and lamina propria) or mesenteric lymph nodes were analyzed using an Attune NxT flow cytometer (Life Technologies, Thermo Fisher Scientific, Waltham, MA). For analysis of immune cell subsets, single cell suspensions were stained with fluorescent antibodies directed against the following lineage markers: CD3, CD4, CD8, CD19, F4/80, c-kit, CD38, CD11b, and CD11c (Biolegend, San Diego, CA). To analyze innate lymphoid cell (ILC) subsets, cell suspensions were stained with antibodies against CD127, IL-12Rb2, CD45, NKp46, and a cocktail of anti-lineage antibodies. The gating strategy used for identification of immune cell subsets by flow cytometry is shown in Supplemental Figure 1 and considered the unique staining characteristics of cells isolated from different tissues (i.e., fat tissues *versus* lymphoid tissues such as mesenteric lymph nodes). The lists of antibodies are provided in Table S1.

### Real-time RT-PCR analysis of mRNA responses in fat tissues

Pieces of fat tissue were excised, flash frozen in dry ice and stored at −80°C. The tissues were then crushed and reduced to powder using a bio-pulverizer before adding 1ml of Trizol (Invitrogen, Carlsbad, CA). cDNAs were synthesized by using Superscript V reverse transcriptase (Invitrogen). Real-time PCR was performed as previously described [39]. Results were analyzed using the MxPro RT-PCR software (Agilent, USA), and data were expressed as relative mRNA expression = 2^−ΔΔCt^ where ΔCt = Ct_unknown_ – Ct_HKG_ and normalized against a house-keeping gene (HKG), β-actin. The list and sequence of primers are provided in Table S2.

### Metabolomics

Metabolite profiles in cecal samples were analyzed by proton nuclear magnetic resonance (NMR). Briefly, intestinal scrapings were resuspended in phosphate buffer (4.3 mM Na_2_HPO_4_*7H_2_O, 1.5 mM KH_2_PO_4_, 2.7 mM KCl), homogenized and filtered as described [40]. Aliquots of prepared extract (70 μl each) were mixed with 19 μl of 9 mM trimethylsilylpropionic-2,2,3,3-d4 acid (TSP) in D2O. The samples were transferred into microcoaxial NMR tube inserts (60 μl volume, 2.02 mm OD, Wilmad Lab Glass, Co.), and then placed inside a 5 mm NMR tube. Proton (1H) NMR spectra were acquired at 25°C using a Varian INOVA operating at 600 MHz (14.1 T) as described previously (47), with an acquisition time of 4 seconds, interpulse delay of 6.55 s, and 1,680 transients (3 h of signal averaging). Spectra were pre-processed using Varian software and were baseline corrected using the Whittaker Smoother algorithm. For multivariate data analysis, spectra were binned to reduce the dimensionality and mitigate peak misalignment [41]. A dynamic programming based adaptive binning technique was employed [42], using a minimum and maximum distance between peaks in a single bin of 0.001 and 0.04 ppm, respectively. Quantification of specific metabolite resonances was accomplished using an interactive spectral deconvolution algorithm in MATLAB [42]. All metabolite peak intensities were corrected for equivalent number of protons and normalized relative TSP signal intensity. Peaks were assigned to specific metabolites following a previously described procedure [40].

### Endomicroscopy images

Fat tissues were analyzed using the Viewnvivo (Scintica, Webster, TX), an in vivo endomicroscopy confocal microscope. Briefly, tissues were stored in formalin after collection from the animals at euthanasia. The tissues were then exposed to aquaflavin, and images were taken at 0-15mm from the surface of the tissue, with each image representing 475×475mm area.

### Statistics

Data are expressed as the mean ± standard deviation. Comparisons between two groups were made through unpaired T-tests. Comparisons between four groups were made through two-way ANOVA, followed by the Tukey’s Multiple Range Test. All statistical analyses were performed with the Prism 9 software (GraphPad Software, La Jolla, CA).

## Results

### Paneth cells regulate intestinal permeability in mice receiving high fat diet

Alteration of gut microbial community (dysbiosis) has been shown to be a driver of obesity [9]. Although Paneth cells are a major source of antimicrobial products [43], the body weight of Paneth cells deficient (Sox9^ΔIEC^) mice fed the normal mouse chow was not different to that of control wild-type mice (Figure 1A). Interestingly, the bacteria load in fecal samples was not different between the two groups (Supplemental Figure 1A). However, significant difference was seen at the phylum level with an increase in unknown or unclassified bacteria in the Sox9^ΔIEC^ mice (Figure 1B). At the phylum level, the Sox9^ΔIEC^ mice exhibited a significantly higher population of Firmicutes and a significantly lower population of Bacteroidetes bacteria when compared to the wild-type mice. (Figure 1B). Differences in commensal bacteria were also noted at the genera level, with Sox9^ΔIEC^ mice exhibiting significantly less Lactobacillus and Butyrivibrio (Figure 1C), two bacteria associated with gastrointestinal health and butyrate production, respectively [44,45]. The number of Alistipes, a bacteria associated with increased gastrointestinal inflammation, was significant increase in in Sox9^ΔIEC^ mice [46] (Figure 1C). Along with changes to the bacteria there were also significant changes in the bacterial metabolites found in the cecum, with there being a significant decrease in both Butyrate and Tryptophan in the Sox9^ΔIEC^ mice (Supplemental Figure 2B).

**Figure 1.**
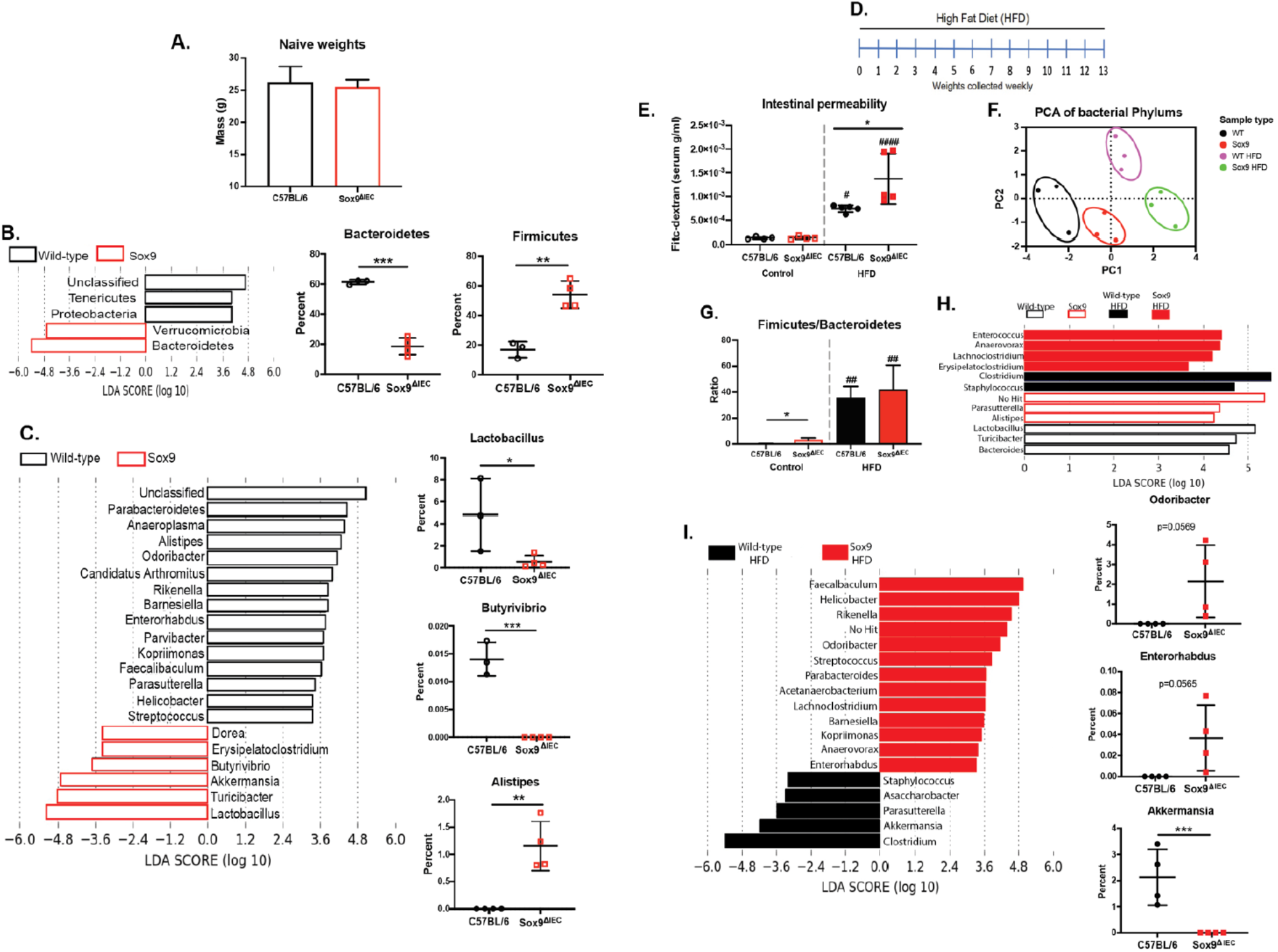
Paneth cells regulate intestinal permeability in mice receiving high fat diet. (A) Weights of the mice before HFD exposure. (B-C) and (F-I) Profile of gut microbiota in Sox9^ΔIEC^ mice. Fecal pellets were collected from wild-type and Sox9^ΔIEC^ mice and microbiota characterized by 16s RNA analysis at (B, F, G) the phylum, and (C, H, I) at the genera level. (D) Schedule of HFD feeding. (E) Intestinal permeability. After 13 weeks of treatment with HFD, mice given FITC-dextran by oral gavage and blood FITC-dextran levels were measured 2 and 4 hours later. (F) Principal component analysis of microbiota phylum of the naïve and HFD groups. (G) Firmicutes to Bacteroidetes ratio between the naive and HFD groups. (H) Microbiota genera in the normal diet and HFD groups. (I) Microbiota genera in fecal samples of wild-type and Sox9^ΔIEC^ mice after HFD. Data represent one of at least three independent experiments with 4-5 mice per group. The alpha value for the factorial Kruskal-Wallis test among classes was 0.05, and the pairwise Wilcoxon test between subclasses. The threshold on the logarithmic LDA score was 2.0. Data are expressed as mean ± standard deviation. Statistical difference between two groups was determined by t-test. Statistical difference between 4 groups was determined by one-way ANOVA multiple comparisons. Statistical significance *p<0.05, **p<0.01, ***p<0.001, and ****p<0.0001 when comparing C57BL/6 and Sox9^ΔIEC^ groups, and #p<0.05, ##p<0.01, ###p<0.001, and ####p<0.0001 when comparing the normal diet to the HFD group.

We next exposed Sox9^ΔIEC^ and control wild-type mice to high fat diet (HFD, 60% calories from fat) (Figure 1D). Because increase in intestinal permeability could drive weight gain and obesity, we analyzed if HFD altered intestinal permeability and the gut microbiota in Sox9^ΔIEC^ mice. The intestinal permeability measured by FITC-dextran showed no difference between wild-type and Sox9^ΔIEC^ mice at steady-state (Figure 1E). HFD increased the intestinal permeability in both groups of mice, and significantly more in Sox9^ΔIEC^ mice than in control wild-type mice (Figure 1E). Principal component analysis shows of the microbiota at the phylum level showed 4 distinct groups, with the mice being fed a normal chow diet being closer together than the mice fed a HFD, we can also see that after HFD exposure the WT (Wild Type) and Sox9^ΔIEC^ mice separate even further (Figure 1F). HFD also altered the microbiota and induced higher Firmicutes/Bacteroidetes ratio in both wild-type and the Sox9^ΔIEC^ mice (Figure 1G). We also found that HFD promotes a significant differences at the genus level, including a decrease in bacteria in the Sox9^ΔIEC^ mice that are associated with gastrointestinal health indicating that the Sox9^ΔIEC^ mice have lost bacteria that have shown to be protective like Aneroplasma and Parabacteroides [47,48] (Figure 1H and Supplemental Figure 2C-E). A more in-depth analysis of the gut microbiota showed differences between Sox9^ΔIEC^ mice and control wild-type mice at the genera levels with increased in Odoribacter and Enterorhabdus, two bacteria associated with increase in inflammation and gastrointestinal distress [49–51]. On the other hand, Akkermansia bacteria, a bacteria associated with mucus production [52,53], was significantly decreased in Sox9^ΔIEC^ mice fed HFD (Figure 1I). Finally, at the genera level HFD-fed wild-type mice showed a trend towards more Lactobacillus and Lactococcus, while Sox9^ΔIEC^ mice trend towards more Faecalibaculum, Helicobacter, and Escherichia (Supplemental Figure 2F).

### Paneth cells deficiency alters gut ILC subsets

In Sox9^ΔIEC^ mice, the dysbiosis alone was not associated with increased intestinal permeability. Thus, we next examined whether lack of Paneth cells affected other parameters of mucosal immune homeostasis. Innate lymphoid cells (ILCs) are among the first immune cells to respond to stimuli from commensal microbes and cytokines produced by epithelial cells [54]. In control wild-type mice, ILC subsets in the lamina propria of consisted of ILC3 followed by ILC2 and negligible (∼1000 cells) number of ILC1 (Figure 2A, Supplemental Table 3). This profile was altered in Sox9^ΔIEC^ mice where ILC2 predominated and ILC3 were significantly reduced (Figure 2A, Supplemental Table 3). We next performed fecal material transplantations to determine whether the profile of ILCs in Sox9^ΔIEC^ mice was driven by the dysbiosis in the absence of Paneth cells. Using naïve conventional WT mice, we gave them 4 doses of fecal material from either a WT or Sox9^ΔIEC^ mouse without antibiotic exposure and analyzed their lamina propria. Lamina propria ILCs in wild-type mice that received fecal material from Sox9^ΔIEC^ mice consisted of a significant increase in ILC2s (Figure 2B). Together, these results suggest that the microbiota of Sox9^ΔIEC^ mice was a major driver of the profile of ILCs in the gut mucosal, and possibly a major regulator of host responses to environment stimuli such as exposure to HFD.

**Figure 2.**
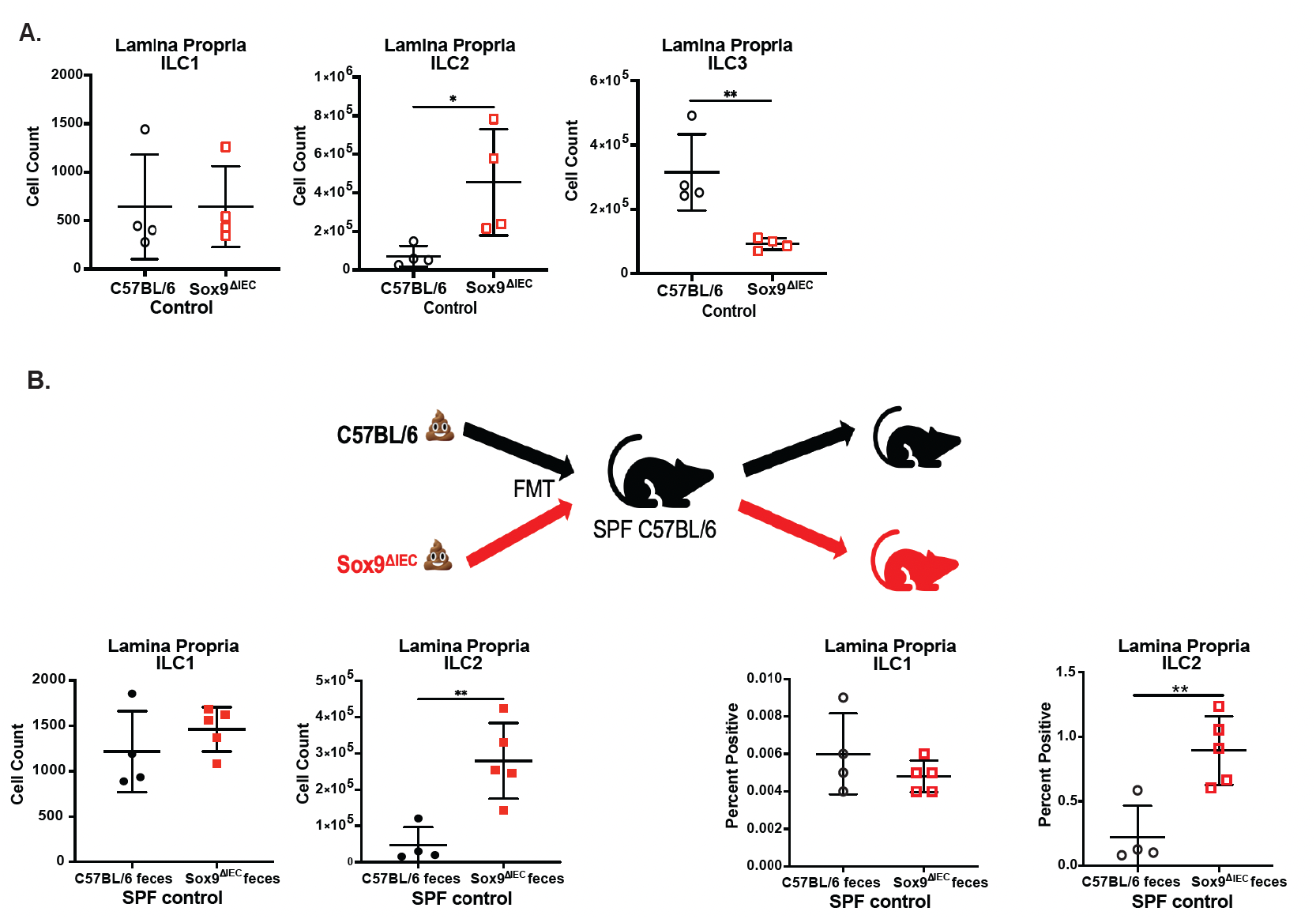
Paneth cells deficiency alters ILC subsets. (A-B) ILC populations (cell number) in the lamina propria before HFD exposure, (B) in wild-type mice given FMTs from Sox9^ΔIEC^ mice. Data are expressed as mean ± standard deviation. Statistical significance *p<0.05, **p<0.01, ***p<0.001, and ****p<0.0001 when comparing C57BL/6 and Sox9^ΔIEC^ groups. Data represent one of at least three independent experiments with 4-5 mice per group.

### Paneth cells deficiency enhances weight gain responses to high-fat diet

Control wild-type and Sox9^ΔIEC^ mice exposed to HFD showed similar levels of food consumption (Figure 3A), but Sox9^ΔIEC^ mice gained more body weight. The difference in body weight was visible by 2 weeks after the beginning of the diet and persisted throughout the 13 weeks (about 3 months) of treatment (Figures 3B). More specifically, Sox9^ΔIEC^ mice accumulated more abdominal fat than control wild-type mice exposed to HFD (Figures 3C-3D). Examination of fat tissues with an endomicroscopic confocal microscope after HFD showed that Sox9^ΔIEC^ mice contained larger adipocytes and more immune cell infiltration (Figure 3D). No difference was noted between brown fat accumulation in wild-type and Sox9^ΔIEC^ mice (Figure 3D). We also analyzed ILCs in abdominal fats since several reports have recently demonstrated their presence in these tissues [55–58]. Although no significant difference was noted between the profile of ILCs in abdominal white fats of mice given HFD, Sox9^ΔIEC^ mice showed trend toward a higher cell count and percentage of ILC1 than in control wild-type mice (Figure 3E). Interestingly, the increased weight gain observed in Sox9^ΔIEC^ mice was associated with switch in the profile of lamina propria ILCs. Thus, HFD significantly increased the cell number of lamina propria ILC1 and reduced the number of ILC2 in Sox9^ΔIEC^ mice (Figure 3F).

**Figure 3.**
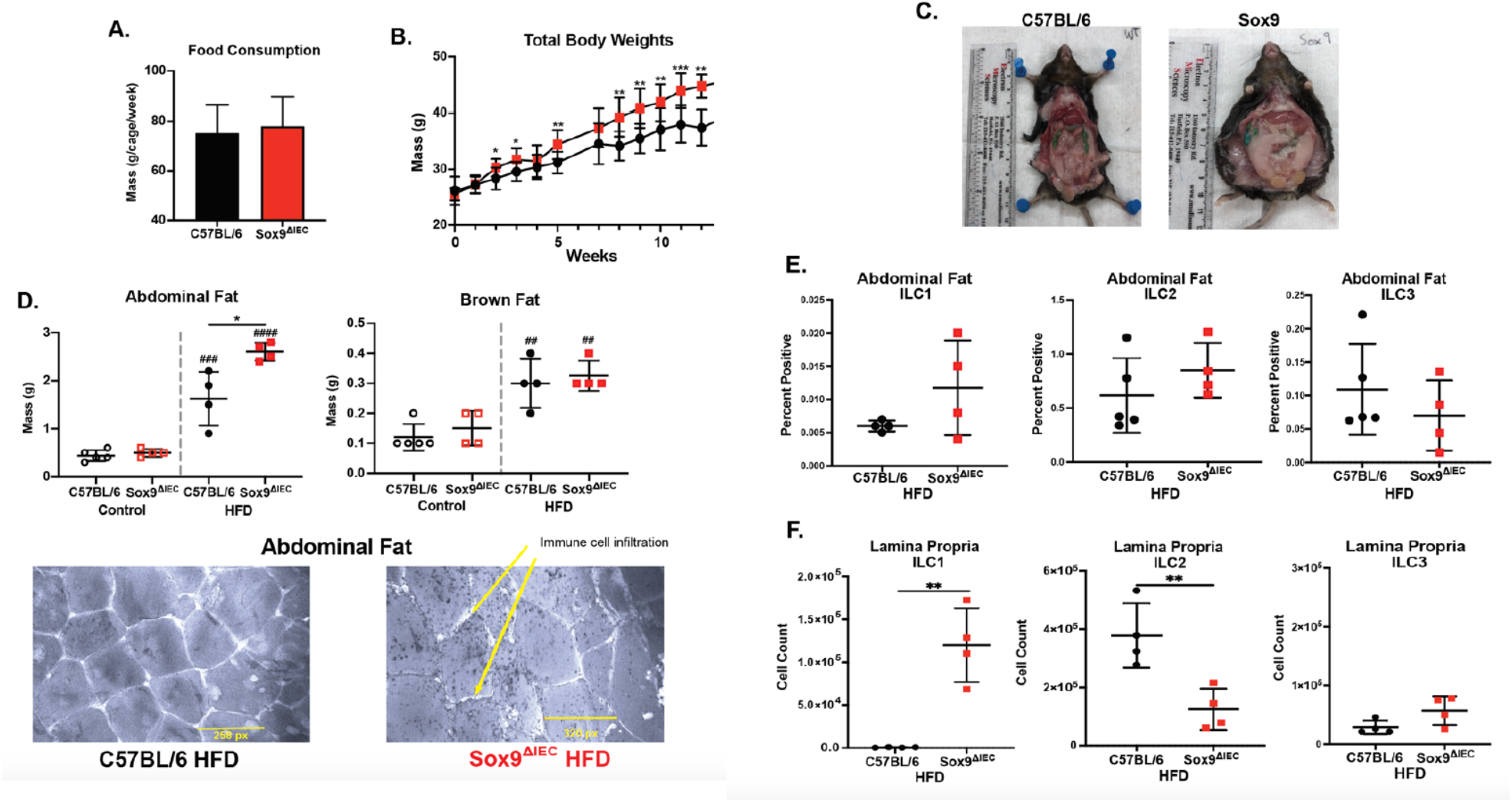
Loss of Paneth cells leads to an increase in weight gain and changes to lamina propria ILC populations after HFD exposure. (A) Food consumption per cage per week during HFD exposure. (B) Weight gain over the 13 weeks of HFD exposure. (C) Images of abdominal fat. (D) Fat weights both before and after HFD exposure. Endomicroscopic confocal microscope images of abdominal fat after HFD. (E) Abdominal fat ILCs after HFD exposure. (F) ILC populations in the lamina propria after HFD exposure. Data are expressed as mean ± standard deviation. Statistical difference between two groups was determined by t-test. Statistical difference between 4 groups was determined by one-way ANOVA multiple comparisons. Statistical significance *p<0.05, **p<0.01, ***p<0.001, and ****p<0.0001 when comparing C57BL/6 and Sox9^ΔIEC^ groups, and #p<0.05, ##p<0.01, ###p<0.001, and ####p<0.0001 when comparing the normal diet to the HFD group. Data represent one of at least three independent experiments with 4-5 mice per group.

### Mice lacking Paneth cells develop more severe metabolic disorders and increased recruitment of myeloid cells and lymphocytes in abdominal fat

In addition to increased size and weight of white fat, signs of obesity include impaired glucose tolerance and metabolic disorder. Sox9^ΔIEC^ mice fed a HFD had baseline blood glucose levels which were higher than those of wild-type mice fed HFD or the control groups (Figure 4A). Following dextrose injection, the Sox9^ΔIEC^ mice show a decrease in glucose tolerance with a higher blood glucose level reached and maintained for a longer period (Figure 4A). We also analyzed expression of leptin and Olr1, which are respectively hormones and lipid receptors known to be important markers in obesity. In this regard, leptin is a hormone produced by adipocytes to induce satiety and stimulate energy expenditure [59] and Olr1 (or LOX1) is an obesity lipid receptor that increases as weight is gained [60]. We found that Sox9^ΔIEC^ mice had significantly decreased expression of *Leptin* mRNA and a significantly increased expression of *Olr1* mRNA in abdominal fat (Figure 4B).

**Figure 4.**
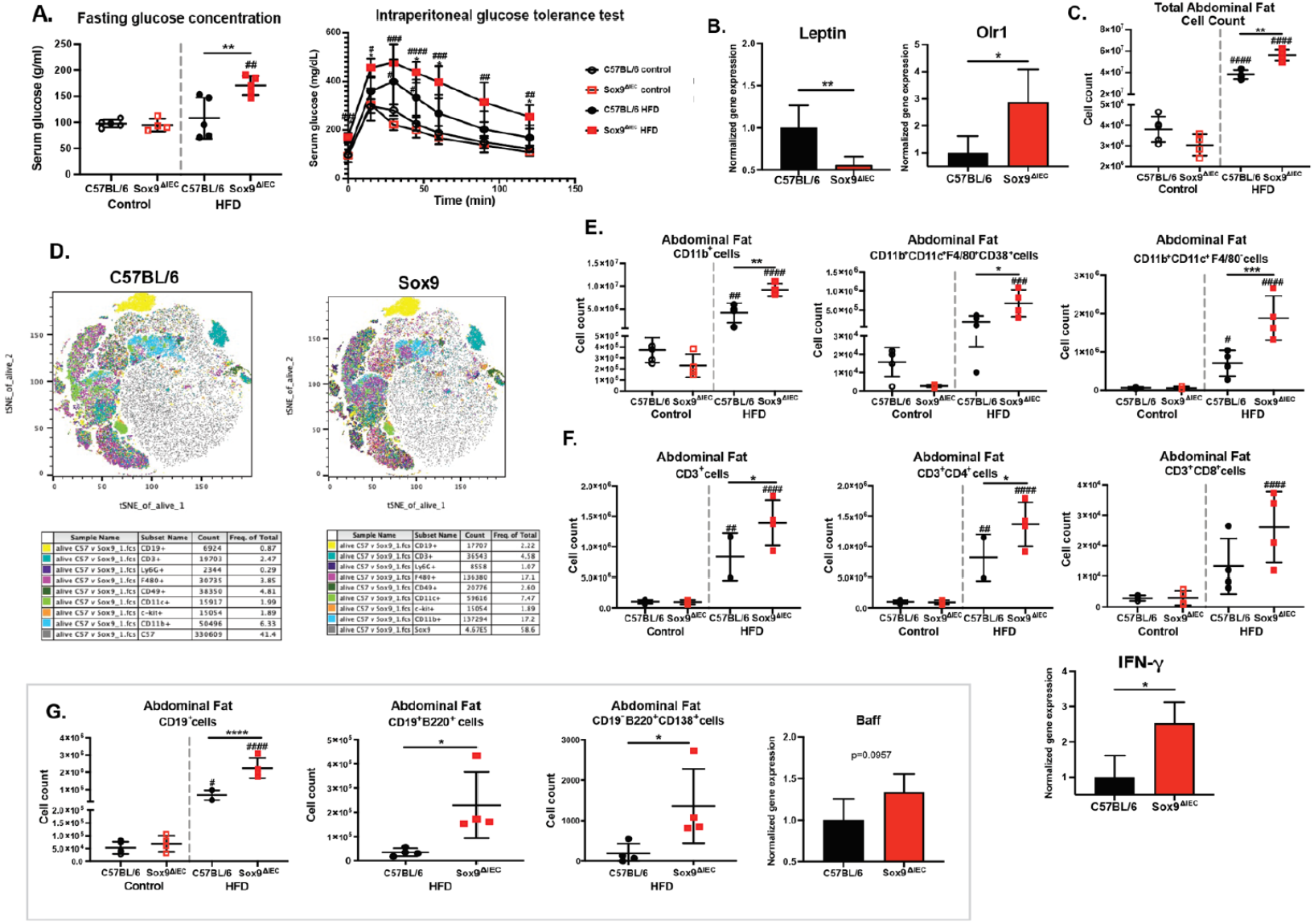
Paneth cell deficiency leads to an increased recruitment of myeloid cells and lymphocytes in abdominal fat after HFD exposure. (A) Intraperitoneal glucose tolerance test before and after HFD. Glucose was injected at 0 minutes and the blood concentration was monitored over 120 minutes. (B) Real-time qRT-PCR analysis of abdominal fat. (C) Total abdominal fat cell counts before and after HFD. (D) t-SNE plot of major cell populations after HFD. (E) Innate immune cell population numbers. (F) T cell population numbers. (G) B cell population numbers. Data represent one of at least three independent experiments with 5 mice per group. The alpha value for the factorial Kruskal-Wallis test among classes was 0.05, and the pairwise Wilcoxon test between subclasses. Data are expressed as mean ± standard deviation. Statistical difference between two groups was determined by t-test. Statistical difference between 4 groups was determined by one-way ANOVA multiple comparisons. Statistical significance *p<0.05, **p<0.01, ***p<0.001, and ****p<0.0001 when comparing C57BL/6 and Sox9^ΔIEC^ groups, and #p<0.05, ##p<0.01, ###p<0.001, and ####p<0.0001 when comparing the normal diet to the HFD group.

Along with changes in metabolic disorders, there is increased evidence that several subsets of immune cells are recruited within the abdominal fat during weight gain [61,62]. Using the gating strategy shown in Supplemental Figure 1, we analyzed the immune cell subsets in abdominal fat before and after HFD exposure (Supplemental Table 4). HFD increased the total number of cells in abdominal fat and significantly more in Sox9^ΔIEC^ mice than in wild-type mice (Figure 4C and Supplemental Figure 3A). Flow cytometry and t-distributed stochastic neighbor embedding (t-SNE) analysis showed higher number of myeloid cells and lymphocytes in the abdominal fat of Sox9^ΔIEC^ mice (Figure 4D). Thus, the significant increase in myeloid cells after HFD correlated with increase in inflammatory macrophages (CD11b^+^, CD11c^+^, F4/80^+^, CD38^+^), dendritic cells (CD11b^+^, CD11c^+^, F4/80^−^) mast cells (CD11b^+^, c-kit^+^), and neutrophils (CD11b^+^, Ly6G^+^) (Figure 4E and Supplemental Figure 3B). The higher number of T cells seen in the fat tissues of Sox9^ΔIEC^ mice after HFD, correlated with significantly higher numbers of T helper cells than in wild-type mice, and significant higher *IFN-γ* mRNA in their abdominal fat tissues (Figure 4F and Supplemental Figure 3C). Sox9^ΔIEC^ mice exposed to HFD had significantly more B cells, and more specifically mature B cells and plasma cells, within their abdominal fat than the wild-type mice (Figure 4G and Supplemental Figure 3C). Finally, the abdominal fat of Sox9^ΔIEC^ mice exposed to HFD contained significantly higher levels of mRNA of chemokines that facilitate inflammatory immune cell trafficking and factors that activate B cell (Supplemental Figure 3D and Figure 4G). Together these results show that intestinal dysbiosis caused by a lack of Paneth cells causes both physiological and immunological changes within the body after exposure to an HFD.

Mesenteric lymph nodes (MLN) are located near the small intestine and are main draining site for immune cell trafficking from gut tissues. Since lack of Paneth cells affected lamina propria ILCs and there were changes within the abdominal fat cell populations after HFD exposure, we examined the profile of the main immune cell subset in MLN. Neither the total number of cells in MLN (Supplementary Figure 3E), nor myeloid cell subsets (macrophages, dendritic cells, mast cells, and neutrophils) were altered in Sox9^ΔIEC^ mice (Supplemental Figure 3F). While HFD reduced the frequency of CD8+ T cells in MLN, there was no difference between control and Sox9^ΔIEC^ mice (Supplemental Figure 3H). Since, lack of Paneth cells did not affect B cells in MLN (Supplemental Figure 3I), these lymph nodes are unlikely involved in connecting changes induced in the small intestinal lamina propria with those seen in the abdominal fat of Sox9^ΔIEC^ mice.

### Fecal materials from mice lacking Paneth cells transfer the obesity-promoting phenotype to germ-free mice

We next performed fecal material transplantation (FMT) into germ-free mice to delineate the relative contribution of the dysbiotic gut microbiota *versus* the immune cells regulatory effect of Paneth cells in the obesity-promoting phenotype seen in Sox9^ΔIEC^ mice. The first FMT induced weight loss in GF mice given fecal materials from either control wild-type or Sox9^ΔIEC^ mice (Figure 5A). This indicates the presence of inflammation and digestional distress associated with first exposures to gastrointestinal bacterial colonization. As seen with SPF mice, germ-free mice recipient of FMT from wild-type feces or Sox9^ΔIEC^ showed no differences in food consumption when exposed to HFD (Figures 5B and 5C). However, solid trends toward more weight gain (Figure 5C) and increased signs of glucose intolerance (i.e., higher fasting blood glucose levels and a higher blood glucose concentration upon glucose injection) (Figure 5D) were seen in germ-free mice recipient of FMT from Sox9^ΔIEC^ mice. On the hand, no difference in intestinal permeability was noted between the germ-free mice recipient of FMT from Sox9^ΔIEC^ or wild-type mice (Figure 5E), suggesting that intestinal permeability was more regulated by the cytokines produced by Paneth cells.

**Figure 5.**
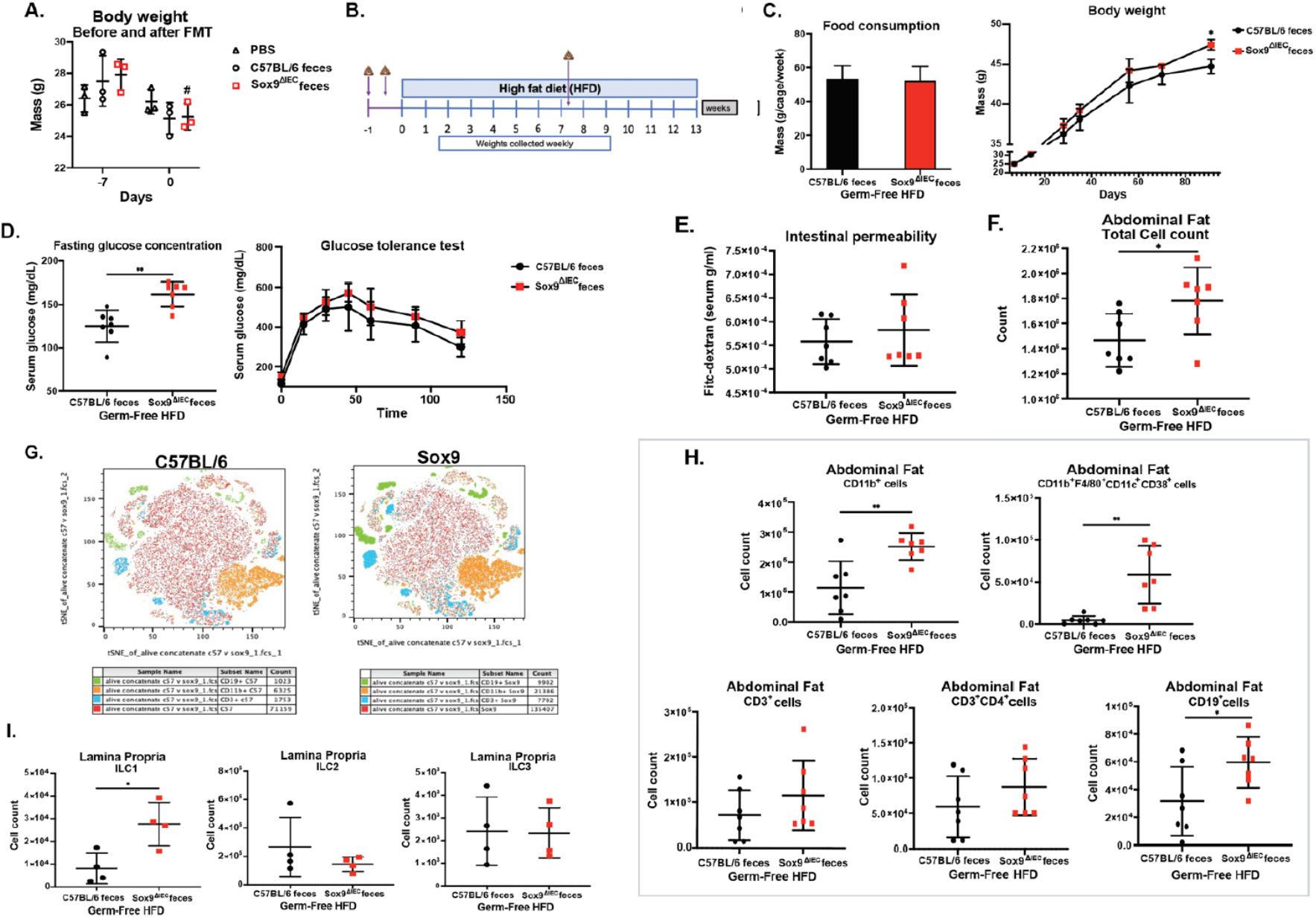
Fecal material from mice lacking Paneth cells transfers a slight increase in physiological and immunological changes to germ-free mice after high fat diet. (A) Weight of the mice before and after 2 FMTs. (B) Schedule of FMT (see arrows) and HFD feeding. (C) Food consumption per cage per week and weight gain over 13 weeks. (D) Fasting glucose concentrations and intraperitoneal glucose tolerance test after HFD. Glucose was injected at 0 minutes and the blood concentration was monitored over 120 minutes. (E) Intestinal permeability. After 13 weeks of HFD, mice were given FITC-dextran by oral gavage and blood FITC-dextran levels were measured 2 hours later. (F) Total immune cell count in abdominal fat. (G) t-SNE plot of major cell populations after HFD. (H) Immune cell subsets in the abdominal fat. (I) ILC populations in the lamina propria after HFD. Data represent one of at least two independent experiments with seven mice per group and are expressed as mean ± standard deviation except in t-SNE plots. The alpha value for the factorial Kruskal-Wallis test among classes was 0.05, and the pairwise Wilcoxon test between subclasses. Statistical difference between two groups was determined by t-test. Statistical difference between 4 groups was determined by one-way ANOVA multiple comparisons. Statistical significance *p<0.05, **p<0.01, ***p<0.001, and ****p<0.0001 when comparing C57BL/6 and Sox9^ΔIEC^ groups.

Analysis of fat tissues of germ-free mice recipient of FMT and exposed to HFD, showed an increase in abdominal fat cell counts (Figure 5F). Flow cytometry and t-SNE analysis of abdominal fat showed an increased infiltration of main immune cell subsets (Figure 5G and Supplemental Figure 4A). More specifically, mice given Sox9^ΔIEC^ feces have more myeloid cells, especially inflammatory macrophages (Figure 5H and Supplemental Figure 4B). We observed no differences in the T cells (Figure 5H and Supplemental Figure 4C). In contrast, we found a significant increase in B cells in the abdominal fat of the mice given Sox9^ΔIEC^ feces (Figure 5H and Supplemental Figure 4C), illustrating the further importance of B cells in abdominal fat, and that the increased amount of B cells is linked to the bacteria found in the Sox9^ΔIEC^ feces.

Finally, analysis of the ILC populations in the lamina propria showed a significant increase in the ILC1s (cell count) in the mice given Sox9^ΔIEC^ feces after HFD when compared to the mice given wild-type feces (Figure 5I and Supplemental Figure 4D). This increase shows the importance of bacteria in driving ILC changes in the gastrointestinal tract in a HFD scenario. We also show that while GF mice were exposed to bacteria and a HFD they still have a lower amount of ILCs in their lamina propria when compared to the wild-type mice (Supplemental Figure 4E). Indicating the importance of bacteria in the development of ILC populations and highlighting that full ILC population could not be replicated in GF mice with 3 fecal material transplants and the presence of an HFD.

It is important to note that in germ-free mice, HFD did not produce the same obesity phenotype as the SPF mice fed the same diet for the same amount of time. We also do not observe as dramatic differences between the groups given Sox9^ΔIEC^ feces *versus* recipients of wild-type feces. Together these findings suggest that most but not all obesity-driving changes that occur in the conventional Sox9^ΔIEC^ mice can be replicated through bacteria given to germ-free mice. Thus, cytokines produced by Paneth cells also play a key role in the protection against obesity.

## Discussion

The prevalence of obesity has steadily increased during the last few decades around the world. Unhealthy diet and alteration of the gut microbial community are among the leading factors known to contribute to obesity [1,2,4,6–8]. However underlying mechanisms are not fully elucidated. Paneth cells are unique intestinal epithelial cells which regulate gut immune homeostasis through production of antimicrobials and cytokines. We now report that Paneth cells play a protective role against obesity. Indeed, dysbiosis in Paneth cell KO mice Paneth cells alone does not lead to obesity. However, it alters the innate immune environment and promotes obesity by enhancing host immunologic and metabolic responses to a high calorie diet.

Alteration of the gut microbial community has been shown to be a driver of obesity [9,63,64]. Consistent with the role of Paneth cells as major producers of anti-microbial products, Sox9^ΔIEC^ mice displayed a dysbiosis of gut the microbial community characterized by an increase in Firmicutes. Interestingly, despite reports linking dysbiosis to weight gain, Sox9^ΔIEC^ mice had unchanged intestinal permeability and failed to spontaneously gain weight. Since dysbiosis can have far reaching consequences throughout the body [65,66], our finding supports the notion that pathologies are not induced by mere dysbiosis but rather loss of specific “protective” bacteria or gain of “pathogenic” bacteria. In this regard, despite the increase in Firmicutes bacteria in the Sox9^ΔIEC^ mice, these gut microbiotas of these mice retain bacteria that are associated with gastrointestinal health such as Anaeroplasma and Parabacteroides [47,48]. Furthermore, the Anaeroplasma and Parabacteroides decreased after exposure of Sox9^ΔIEC^ mice to an HFD confirming their role in health. Thus, despite the dysbiotic state of the gut microbiota of mice lacking Paneth cells, the persistence of protective bacteria prevented weight gain.

Beside their role in the secretion of anti-microbial products, Paneth cells regulate the gut immune homeostasis through secretion of cytokines including IL-17A and TNF-α [22–25]. Our analysis of immune cell subsets in the small intestines of wild-type and Sox9^ΔIEC^ mice showed striking differences in their ILC populations. Innate lymphoid cells are a bridge between the innate and adaptive immune systems and play a vital role in the development of an immune response [54,55]. ILCs can be divided into IFN-γ producing and T-bet-regulated ILC1, the ILC2s that produce IL-5 and IL-13, and require GATA3, and ILC3 which produce IL-17 and/or IL-22 and are dependent on RORγt [67]. It has been shown that the large abundance of ILC3 supports gastrointestinal health through secretion of IL-22, IL-17A/F, and GM-CSF (granulocyte macrophage colony stimulation factor) in the intestinal lamina propria [56–58]. ILC2 can contribute to protection of the intestine against helminth infection and prevent the breakdown of intestinal barrier [68,69]. On the other hand, ILC1s were previously reported to be associated with inflammatory conditions in the gut and in mucosal tissues [70]. We found that lack of Paneth cells in the Sox9^ΔIEC^ mice disrupt the balance of ILC populations, reduces ILC3 while significantly increasing ILC2. Thus, the fact that Sox9^ΔIEC^ mice do not spontaneously develop obesity may be in line with the previous report that ILC2 promotes beiging of white adipose tissue and limit obesity [71]. Microbiota is known to induce the expression of IL-1β and IL-23 which, in turn, activate ILC3 cells [72]. Further, metabolites produced by the commensal microbiota regulate ILCs and tryptophan metabolites, such as indole-3-acetate, can activate ILC3 through stimulation of the Aryl Hydrocarbon Receptor [73]. Tryptophan and butyrate were reduced in Sox9^ΔIEC^ mice further supporting the notion that the role of Paneth is to maintain a microbial microenvironment favorable for expansion of ILC3.

While mice lacking Paneth cells did not spontaneously develop obesity, they were more sensitive to HFD and gained weight faster than their wild-type counterpart and developed more severe obesity and metabolic disorder. The increased ability of Sox9^ΔIEC^ mice to develop obesity upon exposure to HFD is consistent with previous report by other that mice lacking ILC2 or ILC3 cells, but not natural killer cells, are resistant to diet-induced obesity [58]. These authors have also shown that ILC2s isolated from the small intestine, but not ILC2s from white adipose tissue, restores the induction of diet-induced obesity in *Il2rg^−^*^/*−*^*Rag2^−^*^/*−*^mice lacking all lymphocytes [58]. Consistent with the role of ILC2 in obesity, control wild-type mice given HFD showed a switch in the ratio of ILCs with increased number of ILC2. Interestingly, while HFD also switched the ratio of ILCs in Sox9^ΔIEC^ mice, where it induced higher numbers of ILC1 through the body. It has been shown that ILC1 promotes metabolic disorder and other obesity-associated pathologies [74,75]. Thus, reduction of ILC2 and increase of ILC1 could explain the presence of more severe signs of metabolic diseases in Sox9^ΔIEC^ mice. The exact mechanisms which led to reduction/elimination of ILC2 in obese Sox9^ΔIEC^ mice remain to be elucidated. One possibility would be that the most effective utilization of fat in an environment rich in ILC2 has led to excessive proinflammatory polarization of immune signaling in white adipose tissues and this resulted in depletion of ILC2s in fat depot. Indeed, it was shown that TNF-α-induced expression and secretion of the adipokine soluble suppression of tumorigenicity 2 (sST2) leads to depletion of ILC2 [76], possibly through conversion into ILC1. In this regard IL-1β and IL-12 regulate the plasticity of ILC2 and drive their conversion into ILC1 [77,78]. These questions will be addressed in future studies where expression of transcription factors (T-bet, GATA3, RORγt) and other makers (KLRG2 and NKp46) will also be used for a more in-dept analysis of ILCs.

We also made the striking observation that B cells, but not myeloid cells [61,62,79] are the are the major immune cell subset recruited in the fact tissues of obese Sox9^ΔIEC^ mice. This enhances recruitment of B cells was also seen in the fat tissues of germ-free mice that received fecal material from Sox9^ΔIEC^ mice before exposure to HFD. Considering that ILC populations are different in germ-free where the intestines develop in the absence of bacterial pressures [80], these findings highlight the dysbiotic gut microbiome of Paneth cells deficient mice as the initial regulator of the recruitment of B cells. To our knowledge, the role of B cells in obesity remains elusive. However, it is well-known that TNF-induced signaling stimulates the development of lymphoid tissues [81], including Peyer’s patches [82]. More interestingly, it was previously suggested that in atherosclerosis, mouse aorta smooth muscle cells could participate in the formation of tertiary lymphoid tissue via TNFR-1 activation of chemokines which attract lymphocytes and myeloid cells [83]. Future studies will address whether fat tissues of obese Sox9^ΔIEC^ mice develop lymphoid tissues or lymphoid tissue-like structures.

In summary, we have shown that Paneth cells play a protective role against obesity and our findings are consistent with the recent report that administration of Human α-defensin 5 and human β-defensin 2 improve metabolic disorders in mice fed a western diet [84]. We also provide new insights on mechanisms linking Paneth cells and obesity. More specifically, we have shown that the gut community of commensal microbes present in mice lacking Paneth cells stimulates protective mechanisms against obesity which include stimulation of ILC2 throughout the body (see Figure 6). However, upon exposure to HFD, the ILC2 become facilitators of obesity and metabolic disorders possibly through their conversion into ILC1 and thus, further enhancement of inflammatory responses. The function of B cells in adipose tissues remains elusive. But our finding that B cells are the main immune cell subset recruited in the fat tissues of obese Sox9^ΔIEC^ mice provides unique opportunity to address whether fat tissues could be a site for the development of inducible lymphoid structures similar to iBALT induced in the lung after infection [85].

**Figure 6.**
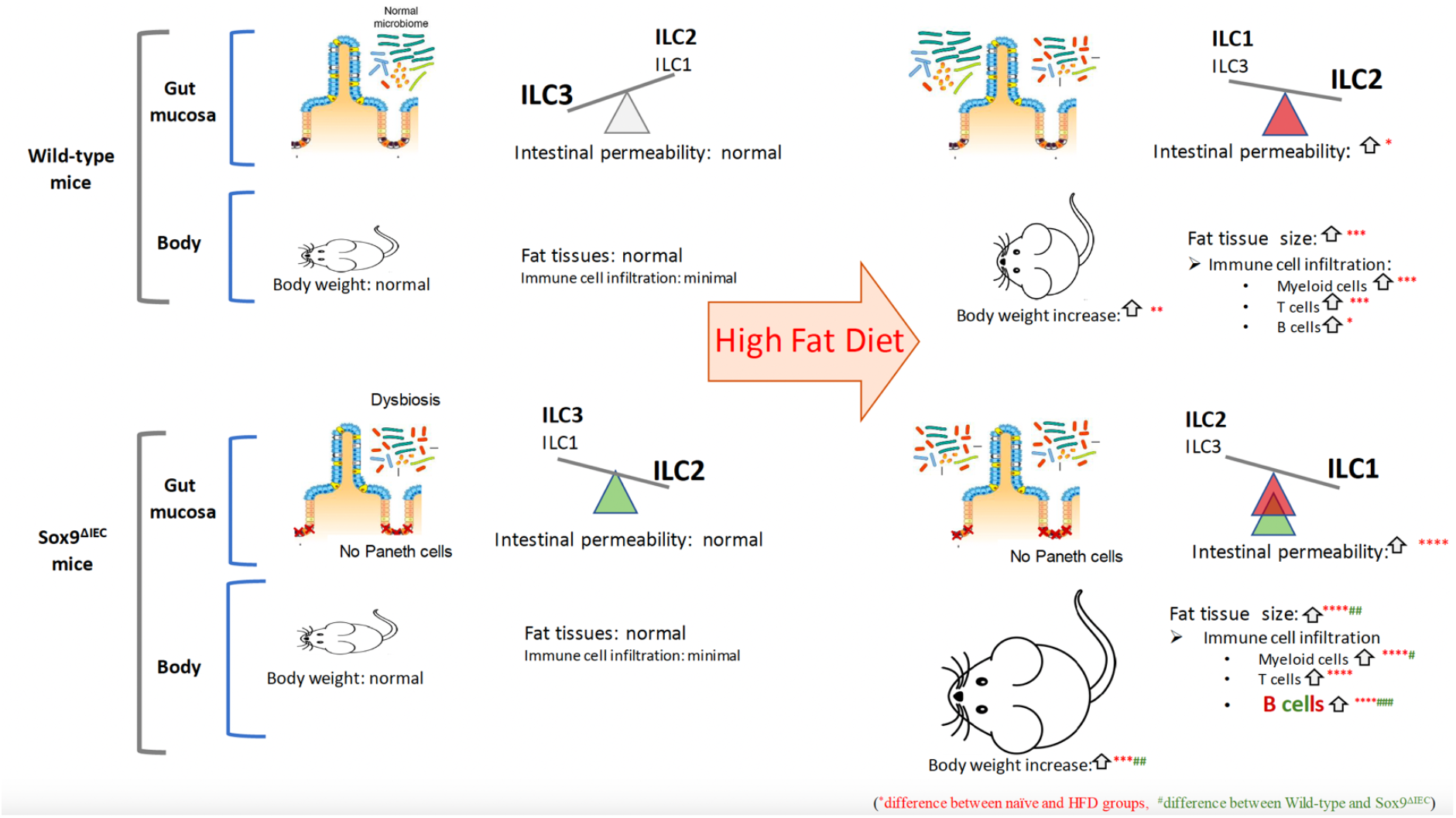
Graphical abstract of Paneth cell regulation of obesity. Dysbiosis alone does not induce obesity or metabolic disorders in mice lacking Paneth cells but alters the ratio of ILCs and increase ILC2. These mice also develop enhanced responses to HFD with greater weight gain and immune cells recruitment in abdominal fat tissues.

## Supplementary Materials

The following supporting information can be downloaded at:,: **Supplemental Figure 1**. Gating strategy for immune cell populations and innate lymphoid cells; **Supplemental Figure 2**. Paneth cells regulate intestinal permeability in mice receiving high fat diet.; **Supplemental Figure 3**. Mice lacking Paneth cells develop more severe metabolic disorders and increased recruitment of myeloid cells and lymphocytes in abdominal fat.; **Supplemental Figure 4**. Fecal material from mice lacking Paneth cells transfers an increase in physiological and immunological changes to germ-free mice after high fat diet; **Supplemental Table 1**. Antibody panels for analysis of immune cells; **Supplemental Table 2**. Primers used for qRT-PCR analysis of mRNA transcripts. **Supplemental Table 3.** Innate lymphoid cell profiles in lamina Propria of WT mice before high-fat diet; **Supplemental Table 4.** Immune cell profiles in abdominal fat of WT mice before high-fat diet.

## Author Contributions

**M.R.J** conducted the experiments, analyzed the data, and wrote the manuscript; **E.K., H.E.S**., Y.J.J., C.P. conducted the experiments and analyzed the data; **W.H., O.Z., and ECB** designed, analyzed the data, and reviewed the manuscript. **P.N.B.** designed the experiments, analyzed the data, and wrote the manuscript.

## Institutional Review Board Statement

Studies were approved by the OSU Institutional Animal Care and Use Committee (IACUC, protocol #2008A0210-R4) and performed on mice over 8 weeks old, in accordance with NIH, OSU IACUC, and ARRIVE guidelines.

## Data Availability Statement

Data is contained within the article and supplementary material are available on request from the corresponding author.

