## Supplemental materials for "Loss of Paneth cells dysregulates gut ILC subsets and enhances weight gain response to high fat diet in a mouse model"

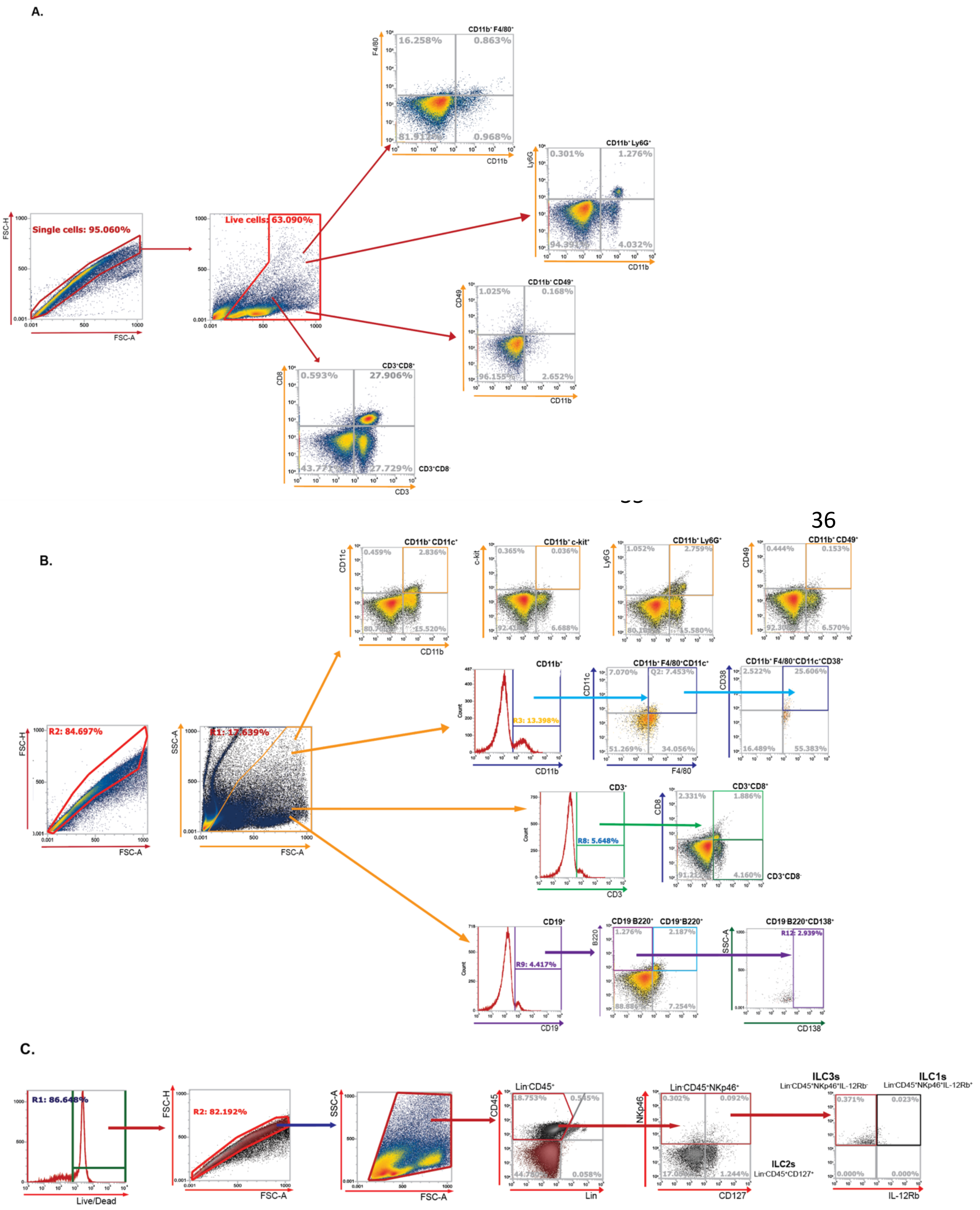

A.
Single cells: 95.060%
FSC-H
FSC-A
Live cells: 63.090%
CD11b+ F4/80+
16.258%
0.863%
0.968%
F4/80
CD11b
CD11b+ Ly6G+
0.301%
1.276%
4.032%
Ly6G
CD11b+ CD49+
1.025%
0.168%
2.652%
CD49
CD3+CD8+
0.593%
27.906%
27.729%
CD3+CD8-
CD8
CD3
B.
R2: 84.697%
SSC-A
CD11b+ CD11c+
0.459%
2.836%
15.520%
CD11c
CD11b+ c-kit+
0.365%
0.036%
6.688%
c-kit
CD11b+ Ly6G+
1.052%
2.759%
15.580%
CD11b+ CD49+
0.444%
0.153%
6.570%
CD11b+
R3: 13.398%
Count
CD11b+ F4/80+CD11c+
7.070%
Q2: 7.453%
51.269%
34.056%
CD11b+ F4/80+CD11c+CD38+
2.522%
25.606%
16.489%
55.383%
CD38
CD3+
R8: 5.648%
CD3+CD8+
2.331%
1.886%
4.160%
CD3+CD8-
CD19+
R9: 4.417%
CD19
CD19-B220+
CD19+B220+
1.276%
2.187%
7.254%
B220
CD19-B220+CD138+
R12: 2.939%
CD138
C.
R1: 86.648%
Live/Dead
R2: 82.192%
Lin-CD45+
18.753%
0.058%
CD45
Lin
Lin-CD45+NKp46+
0.302%
0.092%
1.244%
NKp46
CD127
ILC2s
Lin-CD45+CD127+
ILC3s
Lin-CD45+NKp46+IL-12Rb-
ILC1s
Lin-CD45+NKp46+IL-12Rb+
0.371%
0.023%
0.000%
0.000%
IL-12Rb

**Supplemental Figure 1. Gating strategy for immune cell populations and innate lymphoid cells.** (A) Gating strategy used for flow cytometry analysis of immune cells in mesenteric lymph nodes (MLN). Cell subsets were analyzed after exclusion of duplets and dead cells: macrophages (CD11b$^{+}$F4/80$^{+}$), neutrophils (CD11b$^{+}$Ly6G$^{+}$), NK cells (CD11b$^{+}$CD49b$^{+}$), T helper cells (CD3$^{+}$CD8$^{-}$) and cytotoxic T cells(CD3$^{+}$CD8$^{+}$). (B) Gating strategy used for characterization of immune cell populations in abdominal fat tissues. Cell subsets were analyzed after exclusion of duplets and dead cells: dendritic cells (CD11b$^{+}$CD11c$^{+}$), mast cells (CD11b$^{+}$c-kit$^{+}$), neutrophils (CD11b$^{+}$Ly6G$^{+}$), NK cells (CD11b$^{+}$CD49b$^{+}$), macrophages (CD11b$^{+}$F4/80$^{+}$CD11c$^{+}$), inflammatory macrophages (CD11b$^{+}$F4/80$^{+}$CD11c$^{+}$CD38$^{+}$), T helper cells (CD3$^{+}$CD8$^{-}$), cytotoxic T cells (CD3$^{+}$CD8$^{+}$), B cells (CD19$^{+}$) and plasma cells (CD19$^{-}$ B220$^{+}$CD138$^{+}$). (C) Gating strategy for flow cytometry analysis of ILC subsets. Live cells were analyzed after exclusion of duplets and dead cells: ILC1 (Lin$^{-}$CD45$^{+}$NKp46$^{+}$IL-12Rb$^{+}$), ILC2 (Lin$^{-}$CD45$^{+}$CD127$^{+}$), and ILC3 (Lin$^{-}$CD45$^{+}$NKp46$^{+}$IL-12Rb$^{-}$).

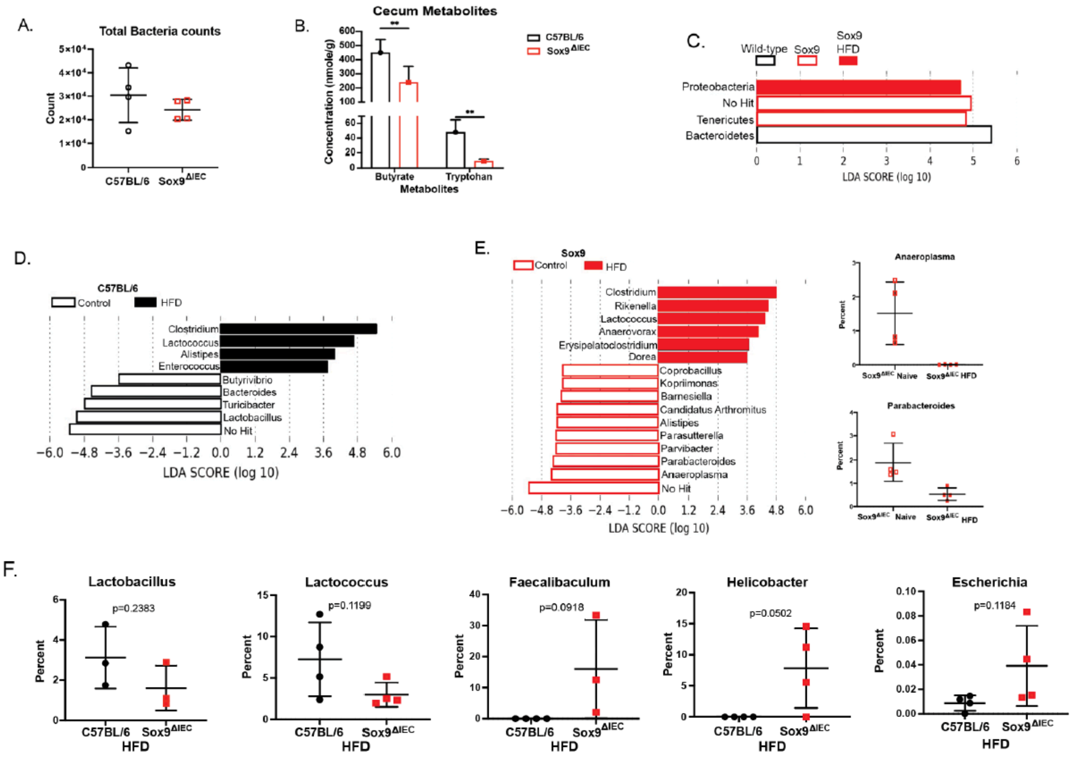


**Supplemental Figure 2. Paneth cells regulate intestinal permeability in mice receiving high fat diet.** (A) Total fecal bacterial counts. (B) Cecum metabolite concentrations, intestinal scrapings were homogenized, and the metabolite profile was analyzed. (C) Profile of phylum differences. (D) Profile of gut microbiota in wild-type mice. (E) Profile of gut microbiota in Sox9$^{\Delta IEC}$ mice. (F) Selected bacterial differences between wild-type and Sox9$^{\Delta IEC}$ mice. Fecal pellets were collected from wild-type and Sox9$^{\Delta IEC}$ mice and microbiota characterized by 16s RNA analysis at the genera level. Data represent one of at least 3 independent experiments with 4 per group. 0.05 was the alpha value for the factorial Kruskal-Wallis test among classes and the pairwise Wilcoxon test between subclasses. The threshold on the logarithmic LDA score was 2.0.

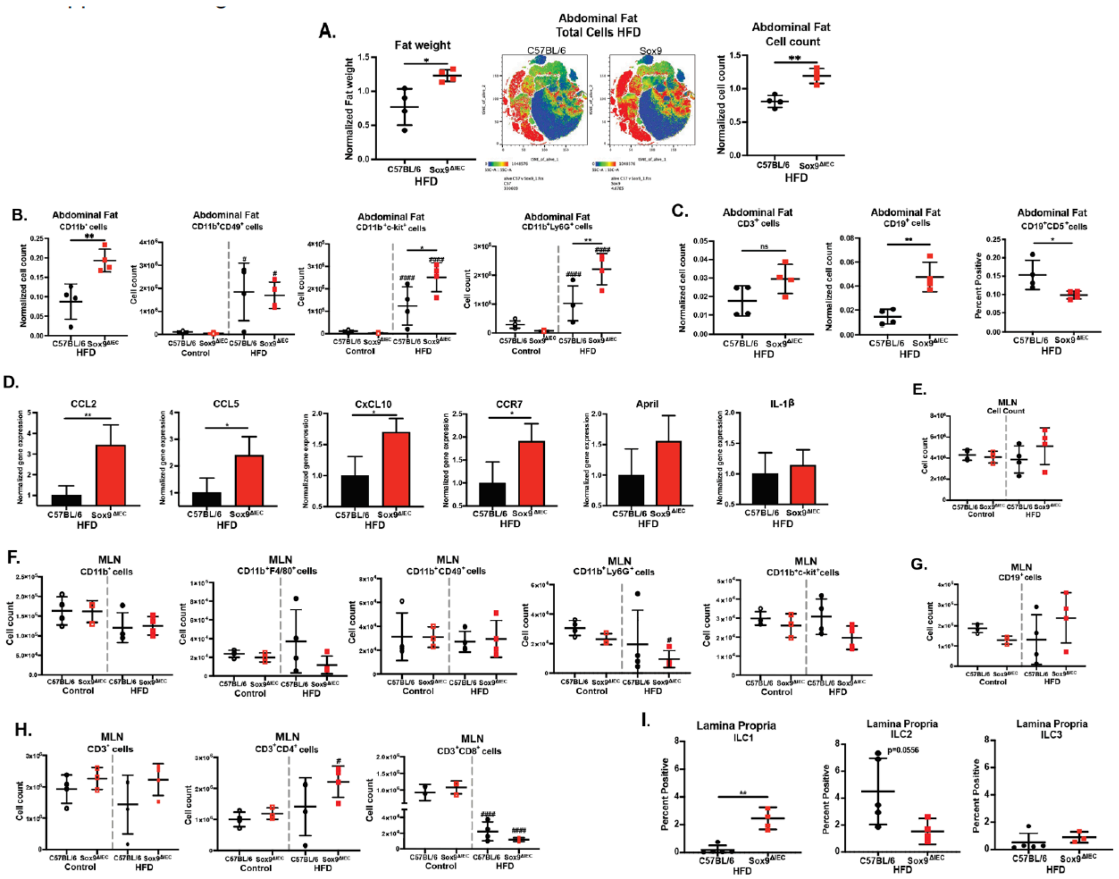


**Supplemental Figure 3. Mice lacking Paneth cells develop more severe metabolic disorders and increased recruitment of myeloid cells and lymphocytes in abdominal fat.** (A) Normalized fat weight, a heatmap using t-SNE mapping showing where the greatest concentration of immune cells is located, and a normalized cell count in the abdominal fat. (B) Normalized to cell count total myeloid cells and other myeloid cell populations observed (C) Normalized to cell count T cells and B cells along with another B cell type. (D) PCR of abdominal fat for cell recruitment markers. (E) Total immune cells in Mesenteric lymph node. (F) Myeloid cell populations in Mesenteric lymph node. (G) B cell population in Mesenteric lymph node. (H) T cell populations in Mesenteric lymph node. (I) Lamina propria ILC percentages after HFD exposure. Data represent one of at least 3 independent experiments with 4 mice in each. 0.05 was the alpha value for the factorial Kruskal-Wallis test among classes and the pairwise Wilcoxon test between subclasses. Data in all bar graphs are expressed as mean ± standard deviation. Statistical difference between two groups was determined by t-test. Statistical difference between 4 groups was determined by one-way ANOVA multiple comparisons. Statistical significance *p<0.05, **p<0.01, ***p<0.001, and ****p<0.0001 when comparing C57BL/6 and Sox9$^{\Delta IEC}$ groups, and . #p<0.05, ##p<0.01, ###p<0.001, and ####p<0.0001 when comparing the naïve group to the HFD groups

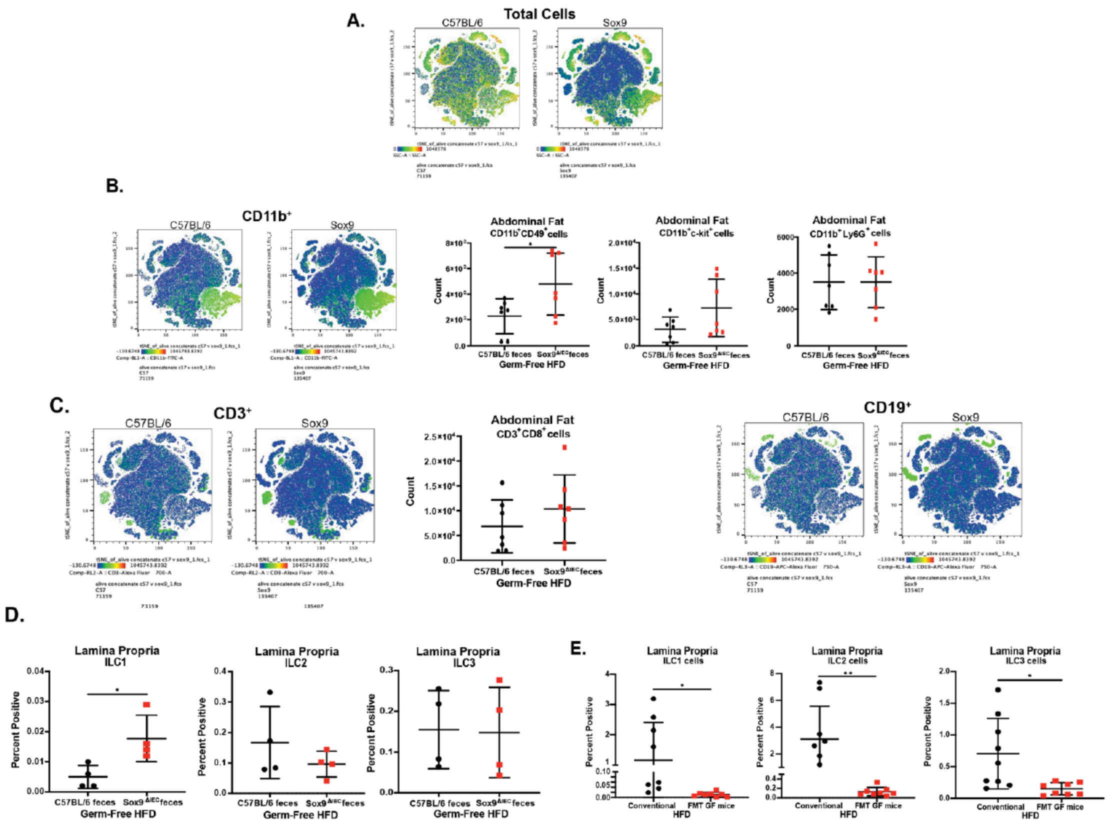


**Supplemental Figure 4. Fecal material from mice lacking Paneth cells transfers an increase in physiological and immunological changes to germ-free mice after high fat diet.**

(A) t-SNE Heat-map immune cell populations. (B) t-SNE Heat-map of myeloid cell populations with breakdown of myeloid cell populations. (C) A heat-map using t-SNE mapping of where T cells and B cells are concentrated. (D) Lamina propria ILC percentages after HFD. (E) The differences in lamina propria ILC between the conventional mice and the GF mice given a fecal material transplant after both have been exposed to an HFD. Data represent one of 2 independent experiments with a combined n of 7 mice. 0.05 was the alpha value for the factorial Kruskal Wallis test among classes and the pairwise Wilcoxon test between subclasses. Data in all bar graphs are expressed as mean ± standard deviation. Statistical difference between two groups was determined by t-test. Significance difference between 4 groups was determined by one-way ANOVA multiple comparisons. Statistical significance $*p<0.05$, $**p<0.01$, $***p<0.001$, and $****p<0.0001$ when comparing C57BL/6 and Sox9$^{\Delta IEC}$ groups.

**Supplemental Table 1. Antibody panels for analysis of immune cells**

Overall immune cell populations

| Laser | Fluorochrome | Marker | Company | Catalog # |
|---|---|---|---|---|
| BL1 | FITC | CD11b | BioLegend | 101205 |
| BL3 | PerCP Cy5.5 | CD8a | BioLegend | 100734 |
| RL1 | APC | c-kit | BioLegend | 105812 |
| RL2 | AF700 | CD3e | BioLegend | 100216 |
| RL3 | APC/Fire 750 | CD19 | BioLegend | 115557 |
| VL1 | BV421 | F4/80 | BioLegend | 123131 |
| VL2 | BV510 | Ly6G | BioLegend | 127633 |
| VL4 | BV650 | CD11c | BioLegend | 117339 |
| YL1 | PE | CD38 | BioLegend | 102708 |
| YL4 | PE Cy7 | CD49b | BioLegend | 108922 |

Brilliant Stain buffer- BD Horizon, BD Biosciences, Catalog # 563794

Innate lymphoid cells (ILCs)

| Laser | Fluorochrome | Marker | Company | Catalog # |
|---|---|---|---|---|
| BL1 | FITC | Lin | BioLegend | 133302 |
| RL1 | APC | IL-12Rb2 | R&D Systems | FAB1959A |
| VL1 | PB | CD45 | BioLegend | 103125 |
| YL1 | PE | CD127 | BioLegend | 135009 |
| YL4 | PE Cy7 | NKp46 | BioLegend | 137618 |

B cell populations

| Laser | Fluorochrome | Marker | Company | Catalog # |
|---|---|---|---|---|
| BL1 | FITC | CCR9 | eBioscience | 11-1991-85 |
| RL1 | APC | CD21/CD35 | BioLegend | 123412 |
| RL2 | AF700 | B220 | BioLegend | 103232 |
| RL3 | APC-Cy7 | CD23 | BioLegend | 101630 |
| VL1 | VioBlue | CD43 | Miltenyi | 130-112-891 |
| VL2 | BV510 | CD5 | BioLegend | 100627 |
| VL3 | BV605 | CD95 | BioLegend | 152612 |
| VL4 | BV650 | CD138 | BioLegend | 142517 |
| YL1 | AF555 | IgA | eBioscience | 12-4204-81 |
| YL2 | PE-Cy5 | IgM | Thermo Fisher | 15-5790-82 |
| YL4 | PE-Cy7 | CD19 | BD Pharmingen | 552854 |

Brilliant stain buffer- BD Horizon, BD Biosciences, Catalog # 563794

**Supplemental Table 2. Primers used for qRT-PCR analysis of mRNA transcripts.**

| Primer | Forward | Reverse | Company | Ref # |
|---|---|---|---|---|
| CCR7 | CTCCTTGTCATTTTCCAGGTGTG | AGTATCACCAGCCCGTTGC | Integrated DNA Technologies (IDT) | 287628855<br>287628856 |
| Leptin | TCACACACGCAGTCGGTATC | GGGTGAAGCCCAGGAATGAA | IDT | 287628857<br>287628858 |
| Muc2 | GAAGCCAGATCCCGAAACCA | GAATCGGTAGACATCGCCGT | IDT | 287628859<br>287628860 |
| CxCL10 | CCAAGTGCTGCCGTCATTTT | AGCTTCCCTATGGCCCTCAT | IDT | 287628861<br>287628862 |
| CxCR3 | ATGCCTCGGACTTTGCCTTT | AAGTCGCTCTCGTTTTCCCC | IDT | 287628863<br>287628864 |
| Ltb4r1 | CGCTCCGAACTATCCCAACA | GGCGAAGGCCAGGATAATGA | IDT | 287628865<br>287628866 |
| Olr1 | CCCTGCTGCTATGACTCTGG | CCTGCTGAGTAAGGTTCGCT | IDT | 287628867<br>287628868 |
| CCR2 | CACCCTGTTTCGCTGTAGGA | CATGGCCTGGTCTAAGTGCT | IDT | 287628869<br>287628870 |

| Primer | Forward | Reverse | Company | Ref # |
|---|---|---|---|---|
| IL-1β | TCGCAGCAGCACATCAACAAG | CCAGCAGGTTATCATCATCATCC | IDT | 143243581<br>143243582 |
| IFN-γ | ACTGGCAAAAGGATGGTGAC | TGAGCTCATTGAATGCTTGG | IDT | 45474120<br>45474121 |
| IL-6 | CCGGAGAGGAGACTTCACAG | TCCACGATTTCCCAGAGAAC | IDT | 53047849<br>53047852 |
| CCR3 | TTGCAGGACTGGCAGCATT | CCATAACGAGGAGAGGAAGAGCTA | IDT | 7554190<br>7554191 |
| April | GGGGAAGGAGTGTCAGAGTG | GCAGGGAGGGTGGGAATAC | IDT | 148065876<br>148065877 |
| Baff | GGACTGATACTGGCGCTGAC | GGGTTTCTGAGGGTACAAA | IDT | 276737123<br>276737122 |
| β-actin | GCGCAAGTACTCTGTGTGGA | GAAAGGGTGTAAAACGCAGC | IDT | 164052399<br>164052400 |
| CxCL13 | TCTCTCCAGGCCACGGTATTCT | ACCATTTGGCACGAGGATTCAC | IDT | 79637639<br>79637640 |

| Primer | Forward | Reverse | Company | Ref # |
|---|---|---|---|---|
| CCL2 | TTCCTCCACCACCATGCAG | CCAGCCGGCAACTGTGA | IDT | 79637641<br>79637642 |
| CCL5 | ATATGGCTCGGACACCACTC | CTTCTTCTCTGGGTTGGCAC | IDT | 45474122<br>45474123 |
| AID | CCAGACTTTGGGTCGTGAAT | TGGCTTGTGATTGCTCAGAC | IDT | 45474106<br>45474107 |

**Supplemental Table 3. Innate lymphoid cells (ILCs) in intestinal lamina propria of WT mice before high-fat diet**

| Lamina Propria | | Total cells (count) | ILC1 (Lin$^-$ CD45$^+$NKp46$^+$IL-12Rb$^+$) | ILC2 (Lin$^-$ CD45$^+$CD127$^+$) | ILC3 (Lin$^-$CD45$^+$NKp46$^+$IL-12Rb$^-$) |
|---|---|---|---|---|---|
| **Experiment 1** (n=4) | Average | 2,154,250 | 95 | 86,079 | 460,487 |
| | STDV | 979,424 | 44 | 4,984 | 29,032 |
| **Experiment 2** (n=4) | Average | 2,654,670 | 121 | 73,672 | 448,562 |
| | STDV | 738,833 | 24 | 2,453 | 34,534 |
| **Experiment 3** (n=4) | Average | 2,393,740 | 200 | 102,034 | 395,383 |
| | STDV | 563,045 | 55 | 1,273 | 21,360 |

**Supplemental Table 4. Immune cell profile in abdominal fat of WT mice before high-fat diet**

| Abdominal fat | | Total cells (count) | CD11b$^+$ | CD3$^+$ | CD19$^+$ |
|---|---|---|---|---|---|
| **Experiment 1** (n=4) | Average | 3,810,000 | 373,049 | 103,960 | 52,854 |
| | STDV | 630,625 | 114,403 | 27,846 | 10,167 |
| **Experiment 2** (n=4) | Average | 3,237,500 | 231,788 | 97,850 | 69,273 |
| | STDV | 525,000 | 105,635 | 33,149 | 3,304 |
| **Experiment 3** (n=4) | Average | 3,150,000 | 315,666 | 103,120 | 35,674 |
| | STDV | 703,562 | 94,729 | 40,152 | 4,770 |
